# Enthalpy-Entropy Compensation

**DOI:** 10.1101/2025.06.10.658876

**Authors:** Philippe Dumas

## Abstract

Enthalpy-Entropy Compensation (EEC) is observed in many unrelated domains. It appears as a strong correlation of the variations ΔΔ*H* and ΔΔ*S* of enthalpy and entropy resulting from experiments performed either at different temperatures on a given system (e.g. the binding of a ligand on a macromolecule), or at the same temperature *T* on related systems (e.g. the binding of related ligands on a macromolecule). In both cases, EEC is characterized by the ‘compensation temperatures’ (ΔΔ*H/*ΔΔ*S*). When a continuous variable *X* (e.g. *X* = *pH*) characterizes the related systems at constant temperature, Θ_*T*_ = (*∂*Δ*H/∂*Δ*S*)_*T*_ may be used in lieu of the ratio of finite variations and when *T* is variable and *X* constant, one may always consider Θ_*X*_ = (*∂*Δ*H/∂*Δ*S*)_*X*_. Thermodynamics trivially imposes Θ_*X*_ ≡ *T*, but also Θ_*T*_ = *T* + *δT* (*X, T*), where the corrective term *δT* (*X, T*) only depends on Δ*G*(*X, T*). The quest for ‘molecular’ explanations of EEC is thus vain: only the value of Θ_*T*_ deserves such explanations. This is illustrated with the denaturation of globular proteins and with the dissociation of hydrophobic peptides from a specific protein. The theoretical estimate Θ_*T*_ = (*T* − *T*\*)*/Ln*(*T /T*\*) with *T*\* *≃* 383 *K* being imposed by experiments, fits well experimental results. Considerations of molecular dynamics (MD) methods led to a theoretical estimate for Θ_*T*_ when no continuous variable *X* exists. One might obtain better MD estimates of Δ*H* and Δ*S* of binding by imposing the correct value of Θ_*T*_.

## Introduction

Enthalpy/Entropy Compensation (EEC) most often appears as a strong linear correlation between the variations ΔΔ*H* and ΔΔ*S* of measured enthalpic and entropic terms in related systems at a given temperature. It was first mentioned, almost one century ago, about data on the solution of different molecules in a given solvent, or of a given molecule in different solvents (1–3). Several examples of EEC, in physical chemistry as well as in biology, with or without water being involved, are shown in SI: enthalpy and entropy of formation of the alcanes (Fig. S1), transfer of alcohols from the gas phase to water (Fig. S2); binding of two molecules in various organic solvents (Figs. S3,S4); formation of RNA helices of various lengths and sequences (Fig. S5); binding of a ligand to a protein at different pHs (Fig. S6). What has to be stressed here is the impressive range of domains having only in common the mention that EEC was observed. The non exhaustive list of 54 different domains reported in (4), those reported in Appendix A in (5), and even the astrophysics field with the thermodynamics of black holes (6), allow everyone to perceive this enormous variety. Theoretical considerations commonly proposed to explain the observed EEC always pertain to the specific example being considered. One is thus faced with a widely, and even universally, observed phenomenon that would require many *ad hoc* explanations, which is quite perplexing.

One goal of this work is of showing that such a quest is unjustified. This is in agreement with one theoretical investigation, among other reports, that showed that EEC is independent of the details of any molecular model and, thus, does not contain any causal link to be uncovered (7). It is worth repeating here an elementary fact supporting this view: that enthalpy and entropy are extensive quantities implies EEC. Indeed, if both Δ*H ≃* Δ*H*_0_ + *n* Δ*H*_1_ and Δ*S ≃* Δ*S*_0_ + *n* Δ*S*_1_, *n* quantifying the size of the system and Δ*H*_0_, Δ*H*_1_, Δ*S*_0_ and Δ*S*_1_ being (almost) constant, then the temperature of compensation ΔΔ*H/*ΔΔ*S ≃* Δ*H*_1_*/*Δ*S*_1_ is (almost) constant, which is the mark of EEC since the variations of Δ*H* and Δ*S* are strongly correlated. This explains what is seen with the alcanes *C*_*n*_*H*_2*n*+2_ with *n* ≥ 2 (Fig. S1), with alcohols of increasing number of carbons (Fig. S2) and with the RNA helices of increasing length (Fig. S5). In the latter case, *n* is the number of base pairs and the occasional inversion of the order dictated by *n* results from the specific effect of the sequence on the thermodynamic parameters (as considered in the many attempts at predicting the melting temperature (8–10)). Of course, EEC is not always the result of enthalpy and entropy being extensive quantities, but it is examined below why it always results from thermodynamics and why only the temperature of compensation requires to be explained at the molecular level. Two examples of determination of the temperature of compensation are given. The often mentioned problem of the lack of significance of EEC, due for example to correlated experimental errors in the Δ*H* and Δ*S* terms (11, 12), will be considered in the conclusion.

### Implications of thermodynamics on EEC

#### EEC at variable temperature and at constant temperature

EEC appears in two different situations that should be clearly distinguished. First, one can consider a given system submitted to the sole variation of temperature. In the chemical/biochemical field, an often encountered example is that of a macromolecule binding a ligand in well defined conditions of pH, ionic strength, etc… Then, the variation of temperature, and only of temperature, will produce a clear EEC, as exemplified in the following. The other situation is when the temperature *T* is kept constant and EEC results from related systems being considered. It is then necessary to introduce a new variable, which will be named *X* in the following, to distinguish the related systems. Obviously, *X* and *T* are independent variables.

#### Introduction of a variable representing related systems

With the example of a macromolecule binding a ligand, the related systems can just differ by a variation of, *e*.*g*. the *pH*: in this case *X* = *pH* is a continuous scalar variable representing completely the different systems. In the study of RNA-helix stability (Fig. S5) (8), the number of base-pairs *n*_*bp*_ is appropriate, even though *X* = *n*_*bp*_ is a discrete variable that cannot account for sequence variability. Finally, the related systems may differ sufficiently that there is no obvious variable to represent each of them. This is common in biochemistry, particularly in pharmaceutical studies, with the study of analogous ligands binding the same target: clearly, a single scalar variable cannot capture the essence of each ligand. It will be seen, however, that characterizing different proteins may nevertheless be achieved, *within the frame of available experimental information*, by a single variable. It turns out to be the case when studying protein stability with *X* = Δ*C*_*p*_ being the variation of heat capacity from the native to the denatured state of each protein. Obviously, one cannot expect that such a fortunate reduction of complexity be often possible.

#### Characterization of EEC with partial derivatives

EEC is commonly characterized by the so-called *compensation temperature T*_*c*_ = (ΔΔ*H/*ΔΔ*S*) calculated with the corresponding variations of Δ*H* and Δ*S*.^1^ When *T* is variable and *X* constant, one can always replace the previous definition of *T*_*c*_ with:

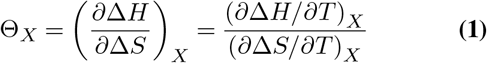

Analogously, when *T* is constant and *X* a continuous variable, one may define:

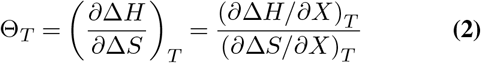

Therefore, two notations Θ_*X*_ and Θ_*T*_ are introduced in lieu of the single notation *T*_*c*_ to differentiate the two situations, a necessity which is sometimes overlooked. Note that, in line with the notations (*∂*Δ*S/∂X*)_*T*_ and (*∂*Δ*S/∂T*)_*X*_, the indices *T* and *X* in Θ_*T*_ and Θ_*X*_ indicate which of the two quantities remains constant. However, with the usual shorthand mathematical notations *∂*_*T*_ Δ*S* and *∂*_*X*_ Δ*S*, they obviously point to the quantity being varied.

#### EEC is always a consequence of thermodynamics: only the value of Θ_*T*_ needs to be explained

We successively consider EEC at variable temperature and at constant temperature. When only the temperature is varied, from the definition of Δ*C*_*P*_ = (*∂*Δ*H/∂T*)_*P*_ and since Δ*C*_*P*_ = *T* (*∂*Δ*S/∂T*)_*P*_, equation 1 implies that, in all situations, the compensation temperature is:

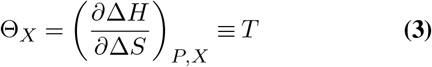

which is nothing else than the ultimate definition of temperature in thermodynamics, irrespective of the system being considered and of any molecular considerations. Therefore, it is worth repeating that such a compensation, and the attached compensation temperature, are trivial consequences of thermodynamics. It is thus unjustified (i) to highlight it as it does not convey any additional information, and (ii) to put forward molecular explanations of it as this is done sometimes (13–16). In fact, this is when EEC is not observed that one should look for explanations. For example, the Δ*H* and Δ*S* obtained by Isothermal Titration Calorimetry (ITC) on a given system at different temperatures should respect Θ_*X*_ *≃* ⟨*T*⟩, where ⟨*T*⟩ is a proper average experimental temperature. Consequently, any significant deviation from the equality should alert either on experimental problems, or on a possibly incorrect model in use.

For EEC at constant temperature, **the only one of interest**, there is a large number of reports stating explicitly, or considering implicitly, that it is definitely not a consequence of thermodynamics, which implies that specific explanations would have to be found. It is worth quoting one very influential paper by Lumry and Rajender: *‘The particular variety of compensation [*…*] is that in which the enthalpy change in an isothermal process is linearly related to the entropy change. This relationship is not in any way a consequence of thermodynamic laws* …*’* (17). A few examples of the quest for a “molecular explanation” of EEC are in (5, 17–21). It is reaffirmed here that this is a false statement and that such a quest is vain. Indeed, when a continuous variable *X* exists, equation 2 joined to Δ*S* = −*∂*_*T*_ Δ*G* implies:

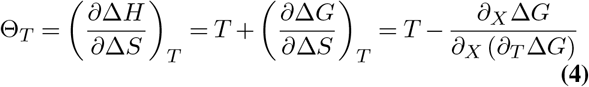

Therefore, Δ*G*(*X, T*) alone determines Θ_*T*_, which implies that *∂*_*X*_ Δ*H* and *∂*_*X*_ Δ*S* **have to be related**. The expression ‘Entropy-Enthalpy Compensation’, which seems to designate a mysterious conjuration producing the compensation, should thus better be replaced with the less connotated ‘correlation of entropy and enthalpy variations’ since thermodynamics shows that this correlation is inescapable. When no continuous variable *X* exists, the partial derivatives should be replaced by the ratio of finite differences:

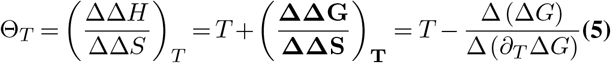

In such a case, an essential limitation to the soundness of Θ_*T*_ is clear: the different systems being considered should be sufficiently similar that (ΔΔ*H/*ΔΔ*S*)_*T*_ approximates a true derivative. This truth is sometimes overlooked, although it was quite clear for Leffler 70 years ago: *‘In a series of related reactions involving* ***moderate*** *changes in structures or solvents the enthalpies and entropies vary, but usually not independently*.*’* (22) (the emphasis on *moderate* is from Leffler). On the contrary, with systems becoming increasingly dissimilar and ultimately unrelated, (**ΔΔG***/***ΔΔS**)_**T**_ in equation 5 would wander more and more erratically between positive and negative values. As a result, Θ_*T*_ would also vary randomly around *T* indicating a poorly defined and meaningless EECC with, in average, Θ_*T*_ ~ *T* (see the conclusion).

Another essential point is that, at variance with the situation of variable temperature for which Θ_*X*_ ≡ *T*, thermodynamics alone does not provide us with any of the partial derivatives or finite differences in equations 4, 5. In conclusion, Θ_*T*_ cannot be obtained without experimental information and, therefore, **only its value, and not EEC, requires molecular explanations**.

#### Another representation of EEC

Representing EEC with (*∂*Δ*H/∂*Δ*S*)_*T*_ = Θ_*T*_ is common, but in no way imposed by thermodynamics. Another possibility is to consider:

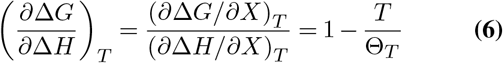

which conveys exactly the same information since the two representations are linked in a trivial way. However, visually, they may be very different. Considering Fig. S4A,B the difference is essentially with the relative variation of the slopes of the two lines appearing in each figure. This is easily understood since, by noting *Y* the slope 1 − *T/*Θ_*T*_, one obtains *dY/Y* = (Θ_*T*_ */T* − 1)^−1^ *d*Θ_*T*_ */*Θ_*T*_. The relative variation of the slope *Y* is thus always greater than that of the slope Θ_*T*_ (as far as T < Θ_*T*_ *<* 2 *T*). The multiplying factor (Θ_*T*_ */T* − 1)^−1^ acts also on the dispersion around the regression lines, which may increase considerably the visual effect. In Fig. S4B (Θ_*T*_ = 374 *K, T* = 303 *K*), the new dispersion is quite moderate because the multiplying factor is 4.3 and the initial dispersion in Fig. S4A is low, but in the inset of Fig. 2B (Θ_*T*_ = 341 *K, T* = 304 *K*), the new dispersion becomes very important because the multiplying factor is 8.2 and the initial dispersion in Fig. 2B is already significant. In no way should this great visual difference be attributed to an independent experimental information. If Θ_*T*_ *< T*, the multiplying factor is negative and the slopes of the two representations are opposite (see the alcanes in Fig. S1). Metaphorically, the representation with (*∂*Δ*G/∂*Δ*H*)_*T*_ provides some kind of a zoom of the representation with (*∂*Δ*H/∂*Δ*S*)_*T*_^2^. This certainly explains why using the Δ*H*,Δ*G* representation was strongly advocated for its ability to give unbiased Θ_*T*_ estimates (23, 24).

#### What can be said about Θ_*T*_ ?

From equation 2, Θ_*T*_ may take positive and negative values since, in the most general situation, the signs of *∂*_*X*_ Δ*H* and *∂*_*X*_ Δ*S* are not bound to be equal. The clearest experimental examples of this are given by vapor-liquid equilibria involving a mixture of two pure compounds, *X* being the molar fraction of one of the compounds (Fig. 1). However, equations 4,5 imply that Θ_*T*_ is always linked to the temperature *T* of the experiment; as a matter of fact, Θ_*T*_ is never far from *T* in the biological field. At constant temperature *T, ∂*_*X*_ Δ*G* = *∂*_*X*_ Δ*H* −*T ∂*_*X*_ Δ*S*, which implies that if *∂*_*X*_ Δ*G* = 0 at a particular value of *X* = *X*_0_, then either Θ_*T*_ = *T*, or both *∂*_*X*_ Δ*S* and *∂*_*X*_ Δ*H* are null. In the latter situation, Θ_*T*_ = 0*/*0 is indeterminate and, by *de l’Hôpital* rule, 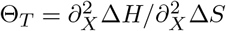. How does this ratio of second derivatives compare with the original ratio of first derivatives? As long as *∂*_*X*_ Δ*H* and *∂*_*X*_ Δ*S* are not null at *X*_0_ where *∂*_*X*_ Δ*G* = 0, Θ_*T*_ = *T* and, by continuity, this is still the case when *∂*_*X*_ Δ*S* and *∂*_*X*_ Δ*H* are null at *X*_0_ (an example of this is given below with the comment following equation 12). Therefore, in all situations, the following necessary and sufficient condition holds:

**Fig. 1.**
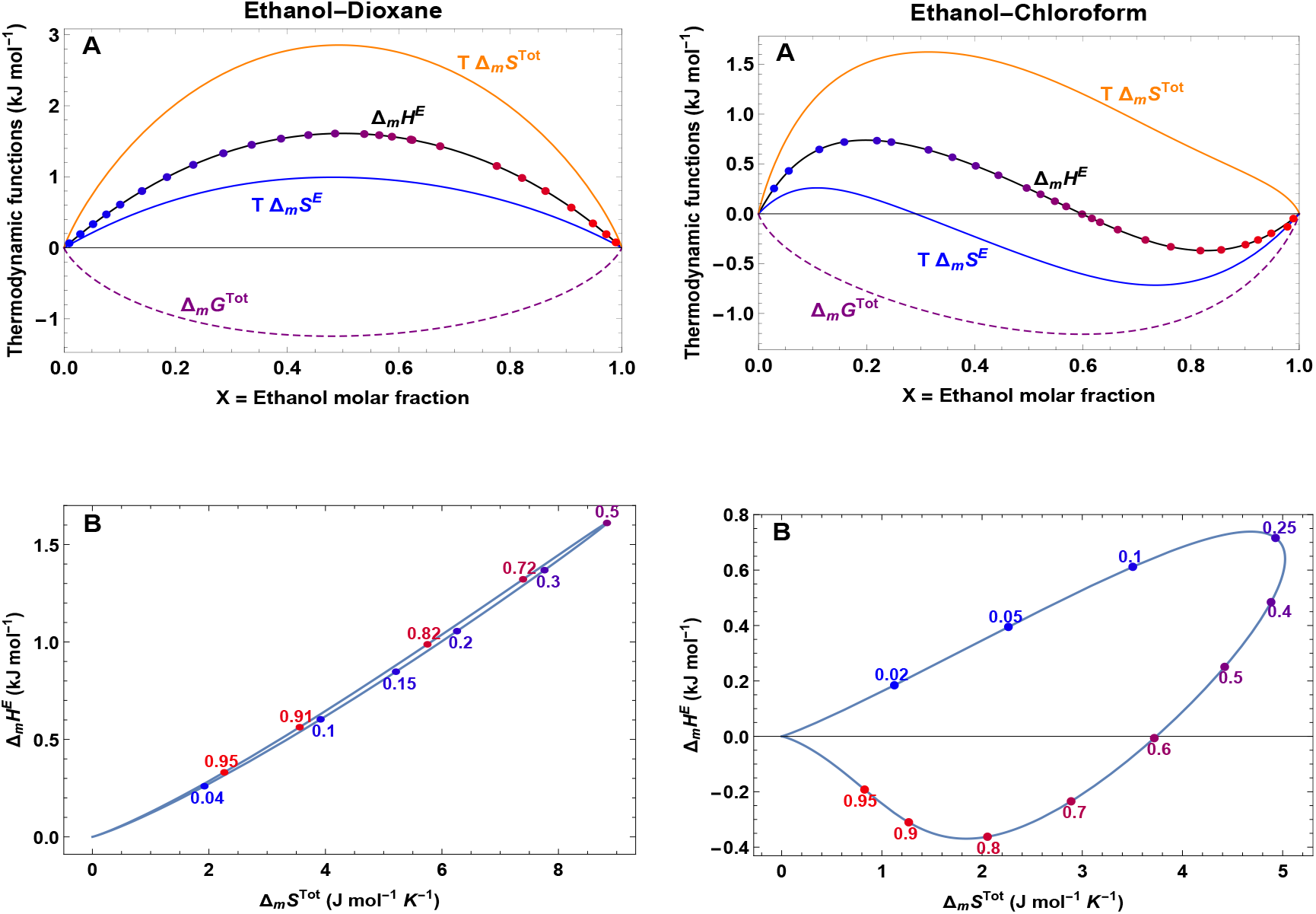
Thermodynamic functions of mixing from vapor-liquid equilibrium and EECC for ethanol-dioxane and ethanol-chloroform. **A** Dependence of the thermodynamic functions of mixing *vs*. the molar fraction *X* (*T* = 323 *K*). The solid curves correspond to semi-empirical equations from (26, 27) and the colored dots to experimental values of the excess enthalpy of mixing Δ_*m*_*H*^*E*^ from (27). The color code varies from blue for *X* = 0, to red for *X* = 1. The total entropy of mixing Δ_*m*_*S*^*T ot*^ is obtained as Δ_*m*_*S*^*Id*^ + Δ_*m*_*S*^*E*^ where Δ_*m*_*S*^*Id*^ = −*R* [*X* ln *X* + (1 − *X*) ln(1 − *X*)] is the entropy of mixing of ideal solutions and Δ_*m*_*S*^*E*^ the excess entropy of mixing. The dashed curves are for the total Gibbs energy of mixing Δ_*m*_*G*^*T ot*^ = Δ_*m*_*H*^*E*^ − *T* Δ_*m*_*S*^*T ot*^ (Δ_*m*_*H*^*E*^ = Δ_*m*_*H*^*T ot*^ since Δ_*m*_*H*^*Id*^ = 0). **B** EECC followed by the point of coordinates (Δ_*m*_*S*^*T ot*^, Δ_*m*_*H*^*E*^) when *X* varies from 0 to 1. Particular values of *X* (chosen for readability) are indicated along the looping curve with the same color code as in **A**. This illustrates (1) that Θ_*T*_, which is equal to the slope of the EECC, may be negative, and (2) that the EECC can show a two-branch pattern with a sharp turning point (ethanol-dioxane at *X ≃* 0.5), exactly as in Fig. S6.

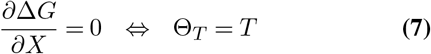

The particular case 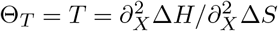 implies that 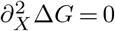 at the particular value *X*_0_. In situations of vaporliquid equilibrium with a mixture of two compounds, this would mean that one is at the limit of liquid-phase separation (25) (p. 258). Therefore, a homogeneous liquid phase in equilibrium with the vapor phase requires that *∂*_*X*_ Δ*H* and *∂*_*X*_ Δ*S* do not vanish at the same value of *X*.

#### Why does the graph of Δ*H* ***vs***. Δ*S* appear so often linear

As shown with the examples in Fig. 2 below and figures S1-S6 in SI, the graph of Δ*H vs*. Δ*S*, that is the Enthalpy-Entropy Compensation Curve (EECC), is often linear, although sometimes with breaks (Fig. S4) and turning point (Fig. S6). This linearity is so common that it is invariably considered as a mark of EEC. However, Fig. 1B for the mixing of ethanol-chloroform shows that, by far, this is not always the case ^3^. Exact linearity is indeed possible as shown in Fig. 2 along with its theoretical interpretation (equation 11) but, in general, the slope Θ_*T*_ of the EECC varies with *X* according to:

**Fig. 2.**
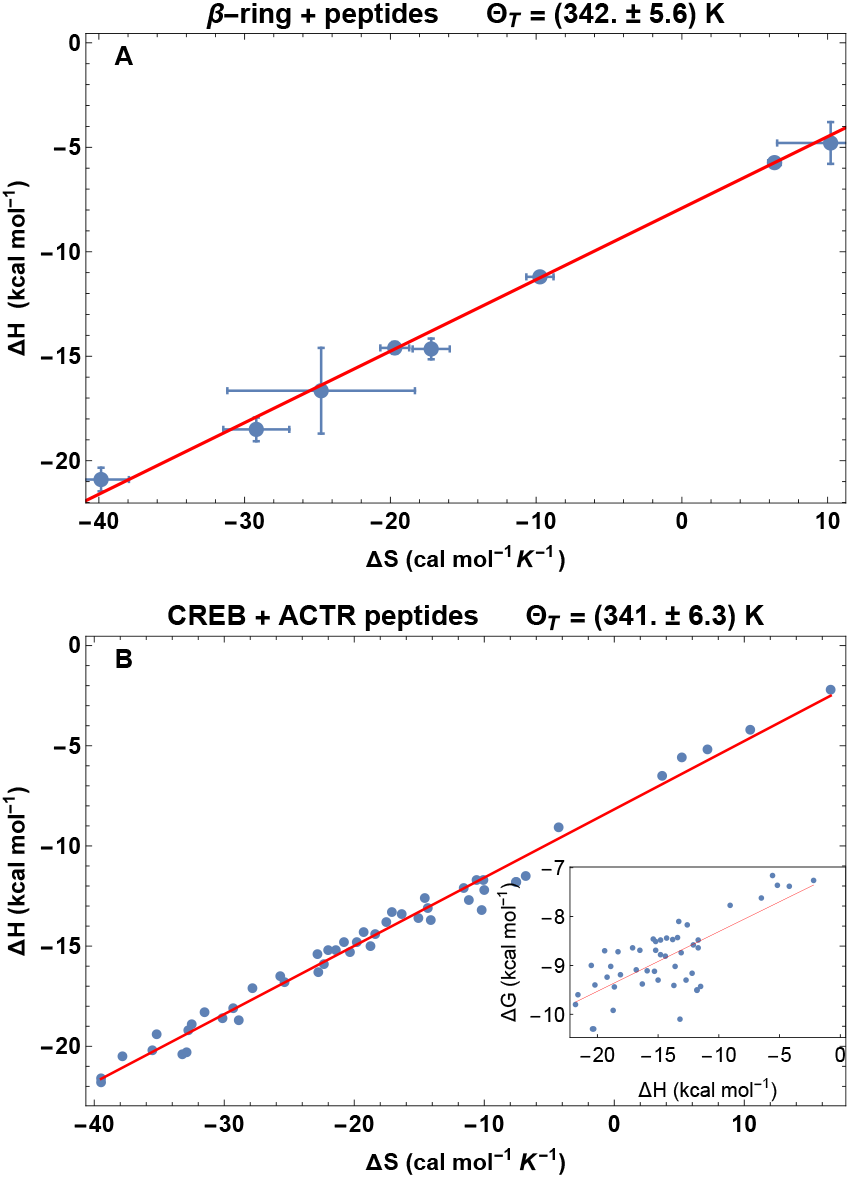
EEC for the binding of related peptides to the *E. coli β*-ring and to CREB-binding protein. The inset for CREB-binding protein shows the EEC representation with Δ*G vs*. Δ*H* (see section *‘Another representation of EEC’*)

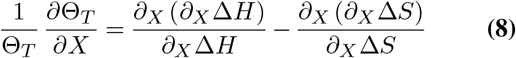

Therefore, if the relative variations of *∂*_*X*_ Δ*H* and *∂*_*X*_ Δ*S* are small, and furthermore of the same sign, then the relative variation of Θ_*T*_ is also small and the EECC is close to be linear. The conclusion is banal: the alleged linearity may simply result from the curvature being masked by experimental errors (if not ignored: see Fig. S2), exactly as for ln *K vs*. 1*/T* in a *van’t Hoff* plot.

### Examples of Θ_T_ estimate

#### EEC and protein stability

##### Context and previous results

Protein stability has been the subject of studies for decades. Experimentally, Differential Scanning Calorimetry (DSC) is the method of choice to obtain thermodynamic information by measuring the variation with temperature of the heat capacity *C*_*p*_ of a protein. Thermal denaturation of the protein is accompanied by a peak in the *C*_*p*_(*T*) curve, which is characteristic of its melting temperature and allows for direct determination of the Δ_*m*_*H* and Δ_*m*_*S* of melting (28). Furthermore, only DSC yields the difference 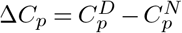 between the heat capacities of the denatured and native states. This difference is invariably positive, which is due (at least in part) to the solvation of hydrophobic compounds as highlighted long time ago by Kauzmann (29). The pH being an important parameter affecting protein stability, DSC experiments by Privalov yielded the variations with the pH of the melting temperature *T*_*m*_ and of Δ_*m*_*H*(*T*_*m*_) for different proteins (30). The results showed that, for each protein, the slope *d*Δ_*m*_*H/dT*_*m*_ was within experimental errors equal to Δ*C*_*p*_ = (*∂*Δ_*m*_*H/∂T*)_*pH*_^4^, itself being independent of temperature within experimental errors. Using normalized values in *cal g*^−1^ for the Δ_*m*_*H*s to allow for their comparison, a linear dependence of Δ_*m*_*H*(*pH*) *vs. T*_*m*_(*pH*) was obtained for each protein and, strikingly, all extrapolated lines concurred to the same point of coordinates 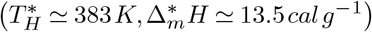 according to:

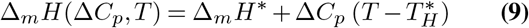

This means that the slope of each line, thus the Δ*C*_*p*_ of each protein, seems to be ‘tuned’ for this convergence to occur. It should be emphasized that this was observed only for *bona fide* globular proteins, and not for other proteins, in particular those with disulfide bonds^5^. This unexpected observation ignited new work, which also revealed a linear dependence of Δ_*m*_*S* (taken at 298 K) *vs*. Δ*C*_*p*_ and a convergence of the Δ_*m*_*S* of all proteins to a common value at 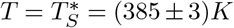(32). This suggested that 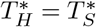, which raised controversy about the interpretation of these observations and on the exactness of the latter equality (19, 33).

##### Results from this work

It is shown here that the latter controversy is resolved since the linear variations of Δ_*m*_*H* and Δ_*m*_*S* with Δ*C*_*p*_ are not independent and, as a consequence, 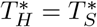 (see SI, part 2). The proof of 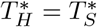 in SI is based on rigorous thermodynamic considerations and, also, on an explicit assumption made in (30): *‘Hence, it seems that we have a right to consider the state of the polypeptide chain near 100* ^0^*C as universal for different proteins’*. In conclusion, assuming this universality, the convergence of all Δ_*m*_*H*(*T*_*m*_) and Δ_*m*_*S*(*T*_*m*_) lines, and the equality 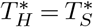, need not any other explanation than that for the convergence of either the Δ_*m*_*H*(*T*_*m*_) or Δ_*m*_*S*(*T*_*m*_) lines. It was recalled in (32) that 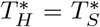 coincides with the temperature for which the Δ*S* of transfer of all liquid hydrocarbons into water is null (34), which is in line with the convergence of the Δ_*m*_*S*(*T*_*m*_) lines.

At this point, one may question the accuracy of the obtained results since both the convergence of the Δ_*m*_*H* lines and of the Δ_*m*_*S* lines appear from lines extrapolated beyond the experimental points that define each of them and *imposing the convergence* yields a reasonable fit of all lines. Since more accurate results would quite probably show approximate convergence, it is suggested that the invoked ‘universality’ is only a useful limiting law as is the ideal-gas law for real gases. Therefore, the equality 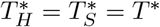 has likely the same validity, or lack of validity, as the invoked ‘universality’.

##### A quantitative estimate of Θ_T_ for protein melting

We are now ready to determine the value of Θ_*T*_ for the EEC observed with the melting of different globular proteins. From the proof of 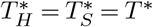 in SI, the dependence of the normalized Δ_*m*_*S* with Δ*C*_*p*_ and *T* is:

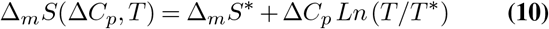

Considering equations 9 and 10, Θ_*T*_ is obtained from equation 2 with *X* = Δ*C*_*p*_ (Δ_*m*_*H*\* and Δ_*m*_*S*\* are constant terms):

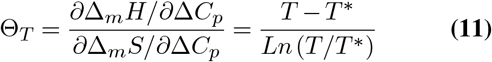

An analogous result was given in (35), but in a wider context and with undetermined parameters. Here, it does not involve any adjustable parameters, but only the experimental value *T*\* = (383 ± 4) *K*. Would 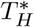 (at the numerator) and 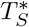 (at the denominator) be different, Θ_*T*_ would be null at 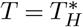 and diverge at 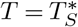, whereas the equality 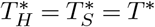 implies that, around *T* = *T*\*, Θ_*T*_ is continuous according to:

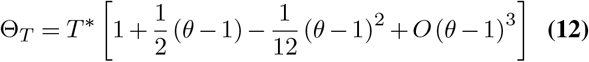

with *θ* = *T/T*\*. Equations 9 and 10 show that *∂*_*X*_ Δ_*m*_*H* = 0 and *∂*_*X*_ Δ_*m*_*S* = 0, and therefore *∂*_*X*_ Δ_*m*_*G* = 0, at *T* = *T*\* (hence at *θ* = 1). Then, as mentioned previously, equation 12 confirms that Θ_*T*_ = *T*\* = *T*, in agreement with equation 7.

##### Experimental verification of Θ_T_ for protein melting

There is a large body of experiments monitoring the stability of proteins. In (19), the results of no less than 3224 experiments were collected, which gave the corresponding values of Δ_*m*_*H* and Δ_*m*_*S* (all at 298 K). This yielded an impressive EEC (99.1 % correlation) with a compensation temperature Θ_*T*_ (exp) = (328 ± 7) *K*^6^. With *T*\* = (383 ± 4) *K* and *T* = 298 *K*, equation 11 yields Θ_*T*_ (theo) = (339 ± 2) *K* and Θ_*T*_ (theo) − Θ_*T*_ (exp) = (11 ± 7.3) *K*. The discrepancy by 1.5 esd is a bit large. Alternatively, if one retains the distinction between 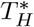 and 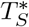 in equation 11 with 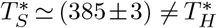, it is obtained Θ_*T*_ (theo) = (332 ± 2) *K* and a discrepancy Θ_*T*_ (theo) − Θ_*T*_ (exp) = (4 ± 7.3) *K* well within the error range. This is not sufficient to conclude firmly that 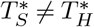.

Beyond protein stability *stricto sensu*, there is a wide interest in peptide/protein interactions which are of considerable biological importance. Such interactions pertain to the previous considerations in the sense that, when the interaction is broken, the peptide is liberated in an unfolded state, which is akin to a ‘denaturation process’. Therefore, if the interaction involves hydrophobic residues, these residues become exposed to the solvent and the previous analysis highlighting the universal temperature *T*\* should apply. This was tested with two series of experiments (Fig. 2). The first one (*T* = 298 *K*) concerns the interaction of the *E*.*coli* DNA-polymerase processivity factor (*β*-ring) with modified peptides derived from the DNA-polymerase stretch interacting with the *β*-ring (36). The second one (*T* = 304 *K*) concerns different methylation patterns of a peptide from the unstructured activation domain of the transcriptional coactivator ACTR interacting with CREB-binding protein (37). In both cases, excellent agreement between Θ_*T*_ (theo) and Θ_*T*_ (exp) is obtained, again without adjusting any free parameter: (339 ± 2) *K* and (342 ± 5.6) *K* for Fig. 2A, and (342 ± 2) *K* and (341 ± 6.3) *K* for Fig. 2B.

##### *Thermodynamic interpretation of* Θ_*T*_

Equation 6 stating (*∂*Δ*G/∂*Δ*H*) = 1 − *T/*Θ_*T*_ yields an immediate thermodynamic interpretation of Θ_*T*_. When Θ_*T*_ *> T*, as for the peptides under consideration, this is the universal expression for the efficiency of a Carnot engine functioning between a hot source at Θ_*T*_ and a cold source at *T*. Since (*∂*Δ*G/∂*Δ*H*)_*T*_ has the same value at a given temperature for all peptides one has (*∂*Δ*G/∂*Δ*H*)_*T*_ = (ΔΔ*G/*ΔΔ*H*)_*T*_, which implies:

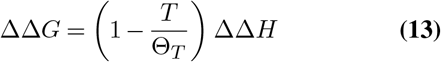

The Carnot efficiency being positive, equation 13 links a negative ΔΔ*H* to the expected decrease in Δ*G* (*i*.*e*. the variation of affinity is enthalpy-driven). With Θ_*T*_ *≃* 341 *K* and *T* = 298 *K*, the decrease is only 1 − 298*/*341 = 12.6% of ΔΔ*H*, and 87.4% is rejected as heat, exactly as a Carnot engine would transform 12.6% of the heat received into work and reject 87.4% into the cold source. Equation 13 can be reformulated in terms of the dissociation constant *K*_*d*_ of a peptide *P* relative to 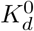 of a peptide *P* ^0^ taken as reference:

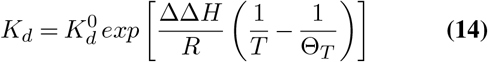

with ΔΔ*H* = Δ*H*(*P*) − Δ*H*(*P*_0_) and *R* the gas constant. This holds for the peptides under consideration with Θ_*T*_ given by equation 11, but this is likely valid for hydrophobic ligands too. Also quite suggestive is the relation ΔΔ*S* = ΔΔ*H/*Θ_*T*_ from the definition of Θ_*T*_ : the variations of Δ*H* and Δ*S* between two ligands are thus linked as in a first-order phase change occurring at Θ_*T*_. Imposing this link might be of interest in molecular dynamics (MD) simulations aimed at evaluating ΔΔ*H* and ΔΔ*S* terms, a notoriously demanding task for which any additional information could be useful. This is examined in the following.

##### Pending test

The agreement of equation 11 with Θ_*T*_ from two series of related peptides at constant temperature is excellent but, according to it, each Θ_*T*_ should also vary with the temperature, which remains to be verified. Equation 12 warns us that a change of temperature by *δT* would only induce a change by *δ*Θ_*T*_ *≃ δT/*2. On one hand, one often cannot increase too much |*δT*| and, on the other hand, Θ_*T*_ is determined with significant uncertainty (see figure 2). A convincing result may not be out of reach with great experimental redundancy.

##### Θ_*T*_ estimate without continuous *X* variable

Here we consider different ligands binding the same target without any suitable variable *X* to characterize each ligand. In such a situation we need to evaluate Θ_*T*_ = (ΔΔ*H/*ΔΔ*S*)_*T*_. It is proposed for that to consider molecular dynamics (MD) methods (see the introduction in (38) and (39)). The free energy variation from one state #0 (taken as reference), to another state #1 only involves the difference in potential energy as follows:

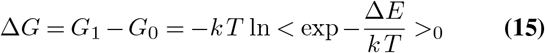

where *k* is the Boltzmann constant, Δ*E* = *E*_1_ − *E*_0_ the variation of potential energy between the two states (a fluctuating quantity during the simulations), and the bracket *<* … *>*_0_ an ensemble average over configurations sampled from the reference state (see (38), chap. 2). There are two possibilities for the states #0 and #1: either they are the free and bound state, respectively, or they are two bound states with two related ligands.

##### Analytical determination of Θ_T_

Normal use of equation 15 requires to estimate the average *<* … *>*_0_ for a specific structure by MD simulations. Instead, in view of obtaining a general result, the average is estimated through analytical calculation following (40) (see also § 2.9.3 in (38)). This is done by considering a sufficiently general probability density function for Δ*E*. Following Amadei *et al*. in (40), a Gamma distribution Γ_1_[*α, β*] depending on two continuous positive parameters *α* and *β* was considered (see SI). Essentially, *α* determines the shape of the distribution, whereas *β* turns out to be a scaling factor that may be taken as 1 without loss of generality. Although it is a versatile distribution since the ‘exponential’, ‘Erlang’ and ‘chi-squared’ distributions are special cases of it (41, 42), its tail for large values of the variable cannot be modified for a given value of *α* (Fig. S7). Since this tail has an enormous importance in the value of *<* exp(− Δ*E/k T*) *>*_0_ for Δ*E <* 0, it is of interest to consider the generalized Gamma distribution Γ_2_[*α, β, γ, µ*] where *γ* allows acting on the tail for large values of the variable (*γ* = 1 and *µ* = 0 result in Γ_2_[*α, β*, 1, 0] ≡ Γ_1_[*α, β*], see Fig. S7A in SI). However, the latter generalized distribution does not yield an analytical result in closed form and one has to rely on a numerical integration to obtain Δ*G* from equation 15 (see SI).

Here, as an example, an exact calculation is made with the simple Gamma distribution Γ_1_[*α, β*]. Since the mean of a variable *x* following this distribution is 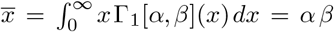, the mean of Δ*E* supposed to follow the same distribution verifies the identity 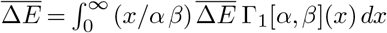. Comparing this with the definition 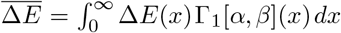 shows that 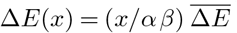 and equation 15 reads^7^:

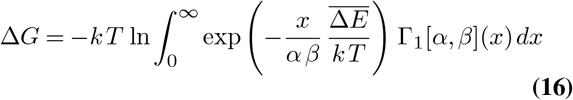

By integration with *Mathematica* it was obtained (per mole):

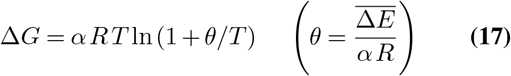

where *θ* has the dimension of a temperature. As mentioned previously, *β* is a physically-irrelevant scaling factor and does not appear in the result. From Δ*G* it is derived Δ*S* = *S*_1_ − *S*_0_ = −*∂*Δ*G/∂T* and Δ*H* = *H*_1_ − *H*_0_ = Δ*G* + *T* Δ*S*:

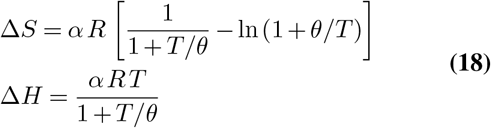

There is an important difference depending on the sign of 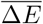: if 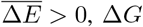 is always defined and positive, but if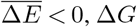 is defined only if 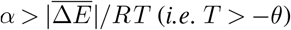 and is negative^8^. Within this crude description of reality, if the reference state #0 is the ligand-free state, *θ* alone characterizes the bound state #1 and, to obtain Θ_*T*_ = ΔΔ*H/*ΔΔ*S*, one should consider two values of *θ*. For closely related ligands, thus for close enough *θ* values, one may consider the approximation:

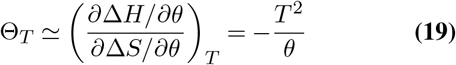

Alternatively, if the reference state #0 is also a ligand-bound state, then ΔΔ*H* = Δ*H* and ΔΔ*S* = Δ*S*, which implies:

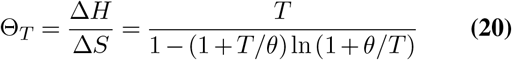

In both cases, Θ_*T*_ and *θ* (and thus Θ_*T*_ and 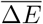), have opposite signs, and Θ_*T*_ → *T* + 0^+^ when 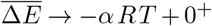 and Θ_*T*_ → ∞ when 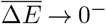. The respective variations of Θ_*T*_ */T* with *T* are represented with the ascending blue curves in figure 3A, which shows that the two situations give quite different results. To be noticed, they give the same value Θ_*T*_ */T* = 1 for *T* = −*θ* (blue dot), that is when Θ_*T*_ from equation 20 becomes undefined due to *T <* −*θ*.

**Fig. 3.**
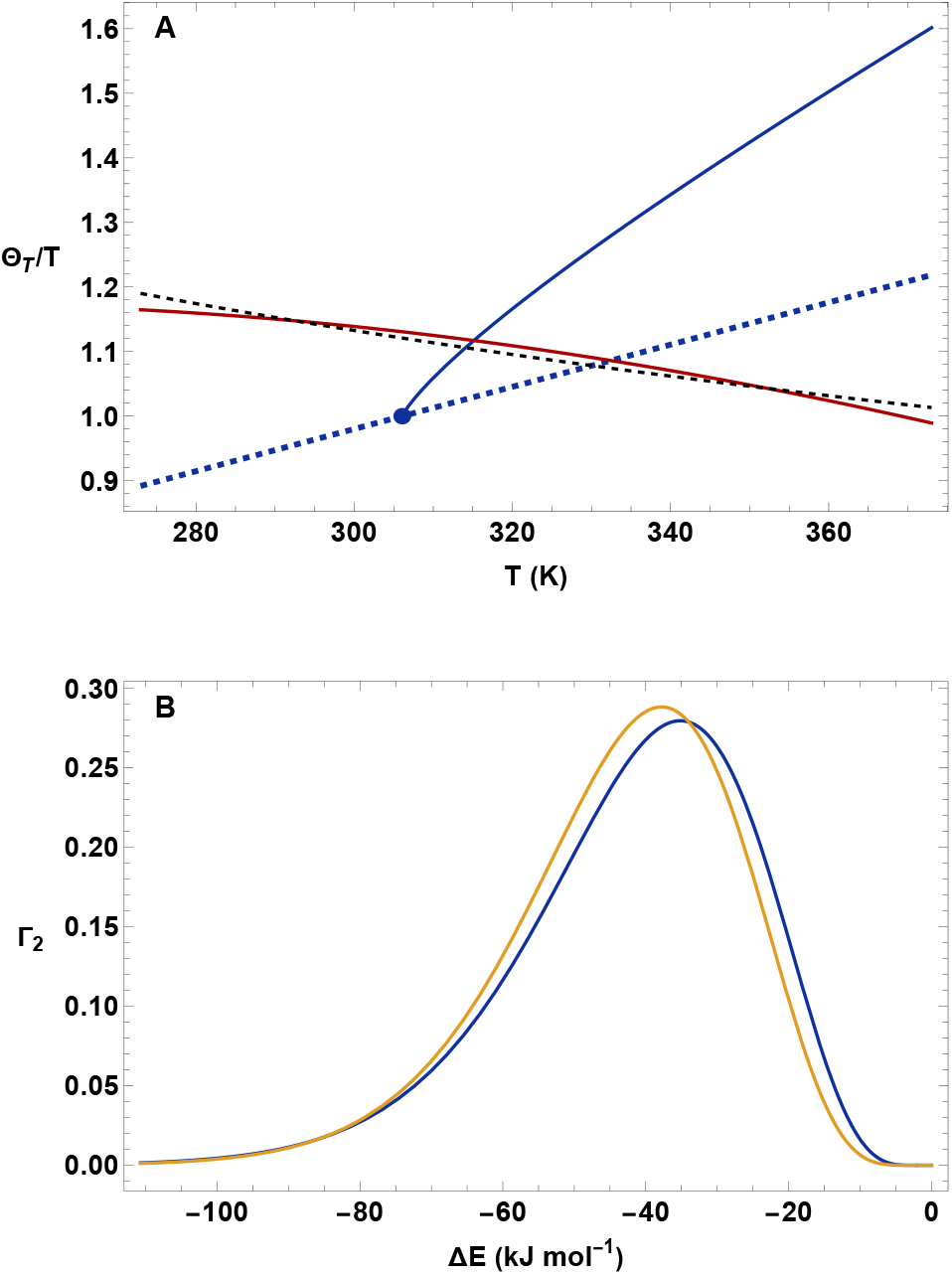
**A**. The Θ_*T*_ */T* curve defined by equation 19 is shown as the dashed blue curve and the Θ_*T*_ */T* curve defined by equation 20 is shown as the solid blue curve (it is undefined for *T <* 306 *K* because the condition 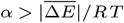 is no more respected). The Θ_*T*_ */T* curve from the Γ_2_ distribution (red curve) is compared to Θ_*T*_ */T* from equation 11 (dashed black curve). The two set of parameters defining the two Γ_2_ curves are given in the following. **B**. Distribution curves Γ_2_ for the two sets of values. The blue curve is defined by 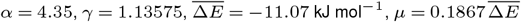, and the yellow curve is defined by 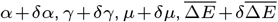 with 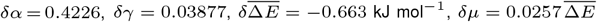(see SI). These values lead at *T* = 293 *K* to Δ*H* = −35.7 kJ mol^−1^, ΔΔ*H* = 5.2 kJ mol^−1^, Δ*S* = −56.3 J mol^−1^ K^−1^, ΔΔ*S* = 15.3 J mol^−1^ K^−1^, *K*_*d*_(#0) = 0.38 mM, *K*_*d*_(#1) = 0.50 mM.

##### Practical usefulness of the Γ_2_ distribution

In order to evaluate the potentiality of Γ_2_, it was attempted to reproduce as closely as possible the evolution with the temperature of Θ_*T*_ */T* for hydrophobic ligands of proteins (equation 11). This is a significant test since the theoretical equation fits well experimental results (Fig. 2). To obtain Θ_*T*_ = ΔΔ*H/*ΔΔ*S*, the values of Δ*H*, Δ*S* for the binding of one ligand (state #0) were determined with a set of parameters 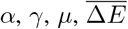, and the values of Δ*H* + ΔΔ*H*, Δ*S* + ΔΔ*S* for the binding of a related ligand (state #1) were determined with the closely related parameters 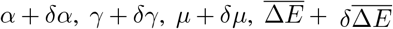. All numerical values and the corresponding Γ_2_ curves are in Fig. 3B (see SI, part 3, about how these values were obtained). Although the result is not perfect, the average slope (if not the curvature) of the dependence of Θ_*T*_ */T vs. T* from equation 11 is well reproduced (Fig. 3A, red and black-dashed curves). The Γ_2_ distribution is thus potentially of practical interest, whereas Γ_1_ is mostly of theoretical interest.

##### Combining Θ_T_ with MD simulations

It is known that MD simulations determine much more accurately the Δ*G* of binding than its Δ*H* and Δ*S* components obtained from error-prone numerical estimates of *∂*_*T*_ Δ*G*. However, it may also be possible to obtain Δ*H* directly and accurately from two MD simulations, one without and one with the ligand being bound (43, 44). Considering two related ligands one would thus obtain either ΔΔ*G* or ΔΔ*H*. In the first case, it would remain to determine ΔΔ*H* and ΔΔ*S* and, in the second case, ΔΔ*G* and ΔΔ*S*. It is clear that knowing the value of Θ_*T*_ would readily allow for their determination without additional MD simulations since one could impose Θ_*T*_ = ΔΔ*H/*ΔΔ*S*. For example, if ΔΔ*G* has been determined, and if Θ_*T*_ ≠ *T* and Θ_*T*_ − *T* is known with enough accuracy, one would readily obtain ΔΔ*S* = ΔΔ*G/*(Θ_*T*_ −*T*) and ΔΔ*H* = ΔΔ*G/*(1 − *T/*Θ_*T*_). This might be tested in many situations since, as already mentioned, Θ_*T*_ from equation 11 should be valid for a series of related hydrophobic ligands, which is common in pharmaceutical studies,

## Conclusions

It is strongly reaffirmed that EEC is inscribed in the foundations of classical thermodynamics, which even ignores the existence of atoms. Therefore, in all occasions, the quest for molecular explanations of EEC is vain. In particular, the common reference to the rôle of water to explain EEC in the biological field (17) is unjustified. Only the value of the compensation temperature Θ_*T*_ requires molecular explanations (which, indeed, may invoke water).

A second conclusion is about the affirmations that EEC is either apparent, or merely an artifact (see (12) for an interesting critical examination of the problem). For example, it is often argued that, when |Δ*G*| is small with respect to |Δ*H*|, Δ*H* and Δ*S* should be correlated with Θ_*T*_ ≈ *T* without the need of invoking EEC (see (7) and references therein). Mathematically, this is true, but the argumentation is implicitly based upon the erroneous conception that EEC requires a physical explanation and, if the need for a physical explanation of EEC disappears, then EEC would not exist. It is then of interest that a purely thermodynamic analysis explained why many EEC observations are expected to show Θ_*T*_ ≈ *T* (45). Moreover, according to equation 7, Θ_*T*_ ≈ *T* results from Δ*G* being not, or little dependent on the variable *X*, which implies nothing against EEC. However, it is exact that gathering too unrelated data respecting |Δ*G*| ≪ |Δ*H*| would produce a completely artefactual EEC with Θ_*T*_ ≈ *T*. One should admit that deciding whether or not a particular situation of this kind corresponds to genuine EEC may not be easy since the dispersion around the regression line is always small. On the contrary, when Θ_*T*_ ≈ *T* without |Δ*G*| ≪ |Δ*H*|, the dispersion around the regression line may become important, which may be considered as a mark of spurious EEC bearing on too unrelated systems (see equation 5 and the comment that follows). A perfect illustration of this was given in (46) with their figure 1 collecting 171 protein-ligand interactions with 32 different proteins and as unrelated ligands as peptides, carbohydrates, nucleotides and synthetic inhibitors. Other examples are in Fig. S1 if alcanes are unduly mixed with other hydrocarbons. A more subtle effect is in Fig. S4 where significant differences within the molecular species are revealed by the need of using two distinct Θ_*T*_ values to explain the data. At the other end, if the dispersion appears to be low with a single Θ_*T*_ value, statistical tests have been proposed to decide whether or not invoking EEC is justified (23, 24). However, these tests only consider Δ*H* and Δ*S* derived indirectly from *van ‘t Hoff* plots for equilibrium data, or Δ*H*^*‡*^ and Δ*S*^*‡*^ from *Arrhenius* plots for kinetic data. These procedures, using experiments at different temperatures to estimate the enthalpic and entropic terms at some averaged temperature, result in correlated errors in both terms (11), whereas their direct determination by ITC or DSC at the desired temperature is much less subject to such errors. Although extremely skeptical about the meaningfulness of EEC when the data are derived from *van ‘t Hoff* or *Arrhenius* plots, Cornish-Bowden recognizes that EEC *‘cannot be easily dismissed as a statistical artefact’* when they are obtained directly by ITC or DSC (47).

A third conclusion is about the explicit determination of Θ_*T*_ through molecular considerations. An example on protein stability illustrates the situation with a continuous variable *X* characterizing sufficiently well the different systems (*i*.*e*. different proteins): the result compared well with different experimental data. Another example when no such variable *X* exists is based upon MD considerations for estimating either the Δ*G* or the Δ*H* upon ligand binding. Although the limitation of the presented approach is obvious since it relies on Gamma distributions to replace the result of an effective MD simulation, it nevertheless illustrates how to estimate Θ_*T*_ without the possibility of using a derivative (*∂*Δ*H/∂*Δ*S*)_*T*_. The suggestion of imposing, when it is known, the value of Θ_*T*_ for a better estimate of ΔΔ*H* and ΔΔ*S* terms from MD simulations requires to be tested.

Finally, it is worth recalling that a continuous EECC may show a turning point (Fig. 1B for ethanol-dioxane and Fig. S6), which corresponds to Δ*H*(*X*) and Δ*S*(*X*) passing through an *extremum* at the same *X*, or at close values of *X*. A discrete EECC may also show a two-branch pattern (Fig. S4). Recognizing this situation is important to avoid obtaining meaningless Θ_*T*_ value. Although up to now widely unnoticed, such patterns for continuous or discrete EECC should not be considered as exceptional and, likely, will be more described in the future.

## ACKNOWLEDGEMENTS

This paper is dedicated to the memory of Peter Privalov (1932-2020) who did so much for our understanding of protein stability. I thank J. Navaza, D. Burnouf, R. Stote, T. Simonson for valuable comments and particularly A. Piñeiro for in-depth discussions. I am indebted to all experimentalists, particularly my colleagues at IGBMC and in E. Ennifar’s team at IBMC.

## Footnotes

1 It is misleading of defining *T*_*c*_ as the ratio of Δ*H* and Δ*S* terms, as done in (5), and not of their variations ΔΔ*H* and ΔΔ*S*.

2 Although one cannot see any other reason for that than pure coincidence, the optical metaphor is astonishingly faithful since the ‘zooming factor’ (Θ_*T*_ */T* − 1)^−1^ compares exactly with the magnifying power (*d/f* − 1)^−1^ of a lens with *d* the object-to-lens distance and *f* the focal length.

3 If the almost linear EECC for ethanol-dioxane is accepted as a *bona fide* example of EEC, then the EEC for the closely related ethanol-chloroform case ought to be accepted too

4 As recalled in (30), this is not at all an obvious result since *d*Δ_*m*_*H* and *dT*_*m*_ in the full derivative *d*Δ_*m*_*H/dT*_*m*_ result from a change of pH, whereas the partial derivative (*∂*Δ_*m*_*H/∂T*)_*pH*_ is obtained at constant pH.

5 The *bona fide* globular proteins were ribonuclease, parvalbumin, lysozyme, *α*-chymotrypsin, *β*-trypsin, cytochrome-c, carbonic anhydrase, and those not showing the convergence were serum albumin, histone H1, ribosomal protein L7 and pancreatic trypsin inhibitor (31).

6 The r.m.s deviation *±*7 *K* was not mentioned in (19) and it was approximately derived from the figure. Same remark for the r.m.s deviation in *T*\* = (383 *±* 4) *K* derived from the figure in (30)

7 The division by *αβ* cannot bear on Γ_1_[*α, β*] which must respect the normalization condition 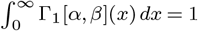.

8 The condition 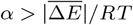 means that the width of the distribution has to increase when 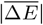 increases. This is not a physical limitation but a limitation of the simple Gamma distribution to represent all situations.

