## Supplemental Text Information for "Enthalpy-Entropy Compensation"

### *Supplementary information for* ‘Enthalpy-Entropy Compensation’

**Philippe Dumas** **** ****

This ‘Supplementary Infomation’ is divided into three parts.
**1** Illustration of ‘Enthalpy-Entropy Compensation’ (EEC) with different examples from the literature.
**2** Determination of $\Theta_T$ related to the problem of protein stability.
**3** Determination of $\Theta_T$ from molecular dynamics calculations (*Mathematica* notebook available on request).

#### 1 Examples of EEC in various fields

Different examples of EEC are shown to give a (small) view of the range of domains where it can be seen. The first two examples are in classical physical chemistry, the third example illustrates the binding of two molecules in various organic solvents and the last two examples are with biological molecules. These examples also illustrate different considerations in the main text, particularly how considering too unrelated data leads to the loss of significant EEC and how an Enthalpy-Entropy Compensation Curve (EECC) may show a *turning point* with an abrupt change of its slope (hence of the compensation temperature $\Theta_T$).

##### Examples of EEC in physical chemistry.

***EEC with the enthalpies and entropies of formation of hydrocarbons from their elements.*** This first example in Fig. S1 is not commonly considered to illustrate EEC because it pertains to enthalpies and entropies of formation of compounds from their elements. It is nevertheless very instructive because it illustrates with the formation of the alcanes $C_nH_{2n+2}$ the clearest possible EEC with $\Theta_T = (215 \pm 4)\,K$ resulting from enthalpy and entropy being extensive quantities. Here, according to equation (8) in the main text, the linearity of the EECC is perfect since the relative variations of the successive $\Delta\Delta H$ terms are equal to the relative variations of the successive $\Delta\Delta S$ terms. Note that, not surprisingly, the linearity breaks down for $n = 1$ since $CH_4$ has no $C-C$ bond to be formed, which is the hallmark of all other alcanes. This also illustrates the comments following equation (5) in the main text by considering other non-saturated hydrocarbons, which leads to a complete loss of significance with a badly defined $\Theta_T = (527 \pm 130)\,K$. The latter value differs by $(229 \pm 130)K$ from the temperature of formation $T = 298K$, which is a large difference in absolute, but only 1.8 times its e.s.d. In contrast, the well-defined $\Theta_T$ of alcanes $(215 \pm 4)\,K$ differs from it by $(83 \pm 4)K$, that is as much as $\sim 21$ times the e.s.d. Therefore, $\Theta_T$ for the alcanes is quite significantly different from $T = 298K$, whereas adding other non-related hydrocarbons leads to a meaningless $\Theta_T$ statistically closer to it.

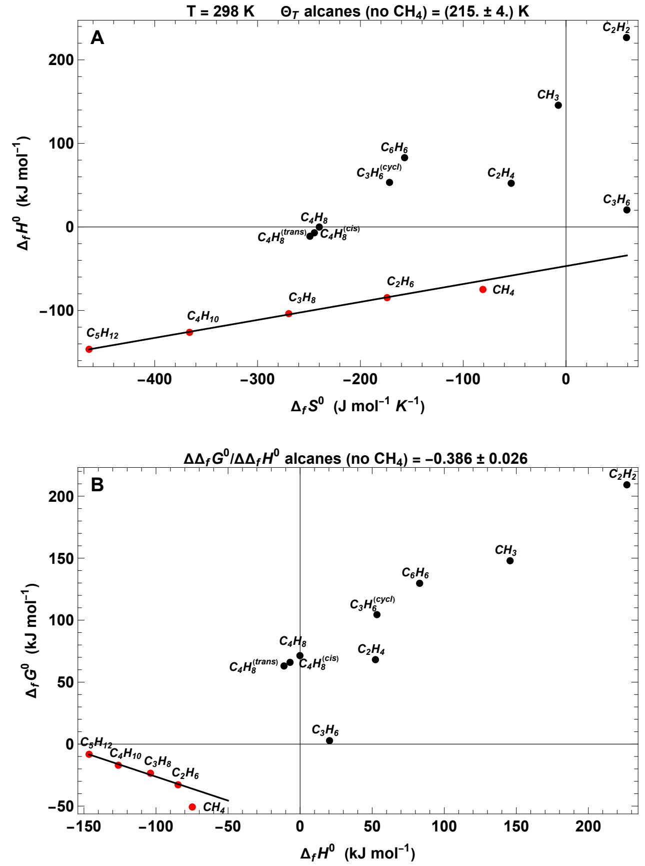


**Fig. S1.** EEC for enthalpies and entropies of formation from their elements of hydrocarbons (taken in the gazeous state at 298 K). Data from Table 2.6 in (1). (A) The figure illustrates the very well defined $\Theta_T$ for the alcanes (if methane is excluded) and the complete loss of significance when other non-saturated hydrocarbons are added. The need of excluding methane is obvious when taking into account that methane has no C-C bond. See also the even more dramatic effect of going from the linear to the cyclic form of $C_3H_6$, which is accompanied with a considerable decrease in $\Delta_f S^0$. (B) Same data represented with $\Delta_f G^0$ vs. $\Delta_f H^0$: note the opposite slopes of the EECC in the two representations due to $\Theta_T < T$ (see section *Another representation of EEC* in the main text).

***EEC with the transfer of alcohols in water.*** Another example of EEC in physical chemistry is with the transfer at 298 K of primary alcohols from the gas phase to water (2). The EECC in Fig. S2 is very well defined. It is apparently linear (black line), but the accuracy is high enough to reveal a small, but significant, curvature justifying a quadratic fit of the experimental points (red curve). Here also this agrees with equation (8) in the main text since the relative variations of the successive $\Delta\Delta H$ terms are now different from the relative variations of the successive $\Delta\Delta S$ terms. As a result, the local slope of the red curve, hence the local compensation temperature $\Theta_T$, varies significantly from methanol to n-pentanol and, therefore, differs significantly from the average linear-fit value $\Theta_T = 265\,K$. This illustrates that, with accurate data, the EECC curvature may be significant even though, by eye, a simple linear fit would seem sufficient.

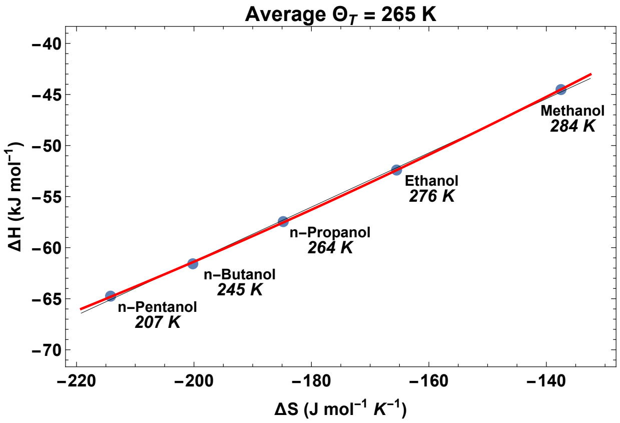


**Fig. S2.** EEC for the transfer at 298 K of primary alcohols from the gas phase to water. Data from (2). The thin black line (differing visually very little from the red curve) results from a linear fit of the experimental points. The red curve is a quadratic fit of the experimental points. The temperatures under each label correspond to the derivative of the quadratic fit, *i.e.* to the local compensation temperature $\Theta_T$.

***The EECC for the effect of various organic solvents on pyrene-cyclophane interaction reveals two distinct groups of solvents.*** The effect of various solvents on the binding of the aromatic four-membered rings pyrene by the cage-like lipophilic molecule cyclophane was studied in (3). The different solvents are shown in Fig. S3.

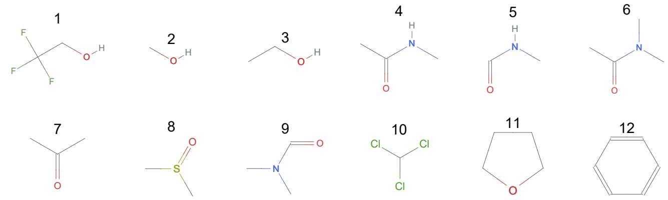

**Fig. S3.** Structure of the different organic solvents used in (3). **1** = 2,2,2-trifluoroethanol, **2** = methanol, **3** = ethanol, **4** = N-methylacetamide, **5** = N-methylformamide, **6** = N,N-dimethylacetamide, **7** = acetone, **8** = dimethyl sulfoxide, **9** = N,N-dimethylformamide, **10** = trichloromethane (chloroform), **11** = tetrahydrofurane, **12** = benzen. This numbering is used in Fig. S4. The figure was made with `ChemicalData` in *Mathematica*.

The resulting EECC in Fig. S4A shows two separate lines with two different compensation temperatures $(373 \pm 2)\,K$ and $(547 \pm 40)\,K$. In fact, the original interpretation in (3) involved a single regression line (dashed line) corresponding to $\Theta_T = (421 \pm 29)\,K$. There are two reasons justifying the splitting into two parts. First, this yields an incomparably better fit, whereas the single dashed line is systematically off the data. Second, the need of doing so is even more obvious when one considers the representation of EEC with $\Delta G$ *vs.* $\Delta H$ in Fig. S4B. This representation conveys exactly the same information as with $\Delta H$ *vs.* $\Delta S$ in Fig. S4A, but increases the relative variation of the slopes of the two parts (see section *'Another representation of EEC'* in the main text). It was recognized in (3) that the $\Delta G$ *vs.* $\Delta H$ graph requires to be split into two part, but this was not extended to the EECC in Fig. S4A as if the two representations would show independent experimental information.

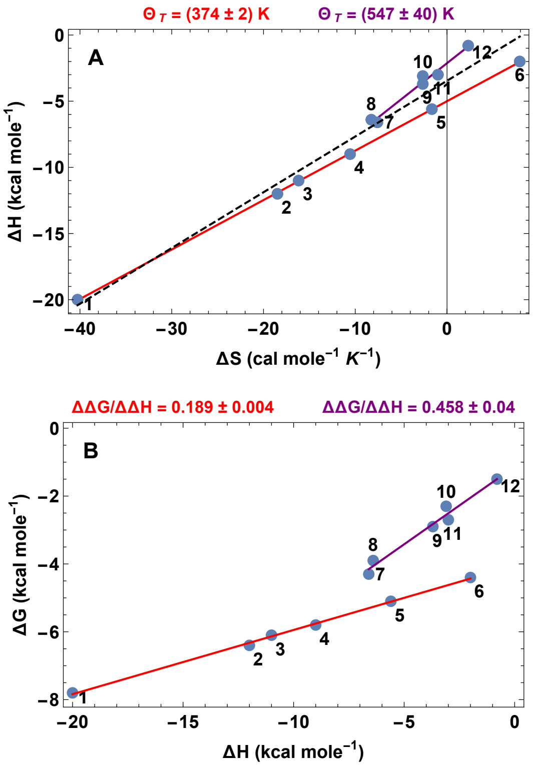


**Fig. S4. A.** EEC for the binding of pyrene to cyclophane in various organic solvents at $T = 303\,K$ (numbering of the solvents as in Fig. S3). Data from from (3). The EECC is split into two parts with two compensation temperatures, whereas the dashed line corresponds to the original interpretation in (3). **B.** Plot of $\Delta G$ *vs.* $\Delta H$ for the same data. Although such a plot does not convey additional information in comparison of A (since B can be deduced exactly from A), it confirms beyond any doubt that the splitting into two parts cannot be avoided.

As far as we are dealing with $\Theta_T$, and not with the existence of EEC, it is legitimate to consider molecular explanations to the splitting of the EECC into two parts. The labelling of the solvents in Fig. S4A shows that one regression line explains those numbered 1 to 6 and the other one explains those numbered 7 to 12. From Fig. S3 we can see that only the solvents 1 to 5 bear a proton that can be involved in H-bonds, and that solvent 6, without such a proton but structurally quite related to solvent 4, is precisely at the junction between the two lines. In addition, apart solvent 9, the solvents 7-12 are perfectly symmetric and have no, or very few (solvent 11), internal degrees of freedom. Interestingly, solvent 9, which seems to be an exception, is in fact a text-book example of 'fluxional molecule' (4), which means that its average conformation is symmetric too like all other solvents 7-12.The firm conclusion based on the observations that the EECC has to be split into two parts thus correlates well with the distinct molecular properties of the solvents. More quantitatively, one may note that the symmetry and the lack of internal degrees of freedom for solvents 7-12 should reduce the overall variation of their entropy of interaction with cyclophane in comparison of solvents 1-6, which is quite clear in Fig. S4A.

**Examples of EEC with biological molecules.**

***RNA helix formation.*** In (5) the thermodynamic parameters at $T = 298\,K$ for the formation of RNA helices of various lengths and sequences were obtained. These data were used in Fig. S5 to illustrate the resulting EEC (not mentioned in (5)). The systematic variation of $\Delta H$ and $\Delta S$ is illustrated by the color code in use. Essentially, the variation follows the number of base pairs, exactly as the variation of $\Delta H$ and $\Delta S$ for the alcohols follows the number of carbon atoms in Fig. S2. Superimposed on this systematic variation is a 'random' variation, which appears with $|\Delta H|$ and $|\Delta S|$ not always increasing with the number of base pairs (see legend of Fig. S5). This 'random' variation corresponds obviously to the influence of the sequence on the stability of the RNA helices.

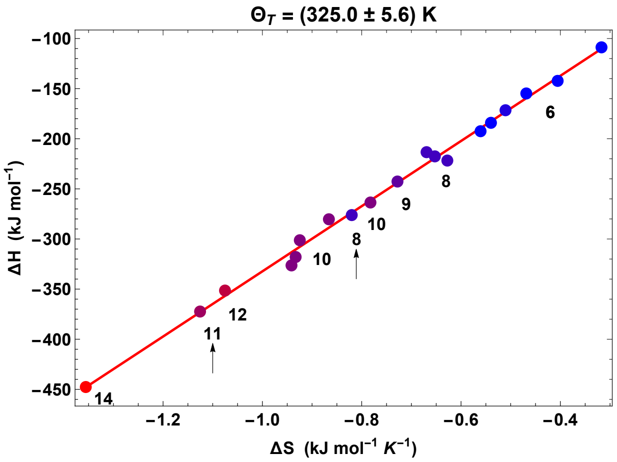


**Fig. S5.** EEC for RNA helix formation at $T = 298K$. Data from (5). The color of each dot represents the helix length and varies from pure blue (6 base pairs) to pure red (14 base pairs. The number of base pairs is indicated to show when the increase of $|\Delta S|$ and $|\Delta H|$ does not follow the increase in the number of base pairs (arrows), which results from the influence of the sequence on RNA stability producing apparently random variations.

***A 'turning point' in the EECC for the interaction of thiocynate with a protein at different pHs.*** In (6) were mentioned and illustrated the results obtained in (7) on the binding of thiocynate to methaemoglobins A at various pHs ($X = pH$) (Fig. S6). These results show a clear 'turning point' on the EECC at $X_0 = pH\,7$ , which was simply noticed in (6). Such a 'turning point' implies that $\partial_X \Delta H$ and $\partial_X \Delta S$ change sign together, which means that both derivatives are null together at $pH = 7$, and thus $\partial_X \Delta G = 0$ too. Therefore, according to equation (7) in the main text, the slope of the EECC at the turning point should be equal to the temperature of the experiment. This is verified as far as it can be by the dashed line in Fig. S6 bisecting exactly the two branches of the EECC.

#### 2 EEC and protein stability

**The linear variation of $\Delta_m H$ *vs.* $\Delta C_p$ implies the linear variation of $\Delta_m S$ *vs.* $\Delta C_p$.** It was noticed empirically in (8) that both the enthalpy and entropy of melting of globular proteins varies linearly with the $\Delta C_p$ of each protein (figure 2 in (8)). This was considered as a significant information on the behavior of these proteins. It is shown here that, in

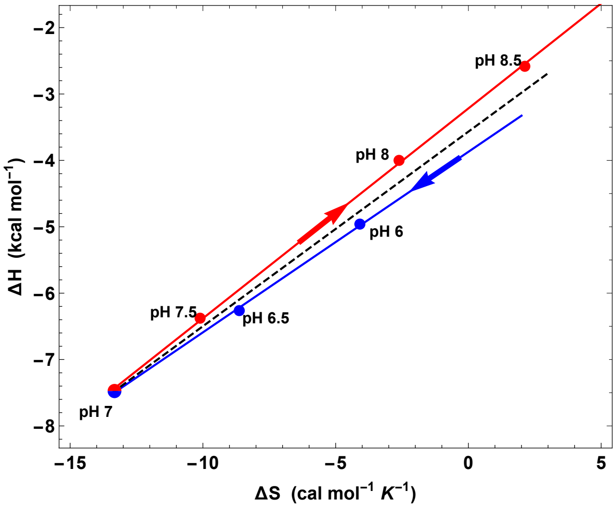


**Fig. S6.** EEC for the binding of thiocyanate to methaemoglobin A at $T = 293K$ and different pHs. Data from (6). The solid lines are linear fits of the experimental points and the arrows indicate the direction of increase of $X = pH$. The *turning point* corresponds to pH 7. The black dashed line indicates the slope equal to the temperature $T = 293\,K$ of the experiment.

fact, the two observations are not independent since any one linearity implies the other one. The following proof derives from classical thermodynamic considerations, along with the explicit assumption made in (8, 9) on the universality of globular-protein melting.

First, it is recalled that the initial observation of Privalov (9) used in Murphy *et al.* (8) is that all $\Delta_m H(T_m)$ of globular proteins converge toward the same value $\Delta_m H^*$ at the temperature $T_H^* \simeq 383\,K$. Now, let us consider one of the two previous empirical observations, for example the linear variation of $\Delta_m H$ with the $\Delta C_P$ of each protein[1]. The only possible relation compatible with both the latter observation and the convergence of all $\Delta_m H(T_m)$ lines is :

$$\Delta_m H(\Delta C_P, T) = \Delta_m H^* + \Delta C_P\,(T - T_H^*) \qquad \textbf{(1)}$$

Then, calculating the partial derivative with respect to $T$ of both members of the latter equation, one obtains $\Delta C_P = \Delta C_P + (T - T_H^*)\,\partial \Delta C_P / \partial T$ (since $\Delta_m H^*$ is a constant term), which imposes $\partial \Delta C_P / \partial T = 0$. Since it was observed empirically that $\Delta C_P$ does not vary with the temperature, equation 1 accounts for all experimental observations relative to $\Delta_m H$. The independence of $\Delta C_P$ *vs.* $T$ readily allows integrating the differential relation $\partial \Delta_m S / \partial T = \Delta C_P / T$, which gives:

$$\Delta_m S = \Delta_m S^* + \Delta C_P\, Ln \frac{T}{T_S^*} \qquad \textbf{(2)}$$

where $T_S^*$ is an arbitrary temperature common to all proteins and $\Delta_m S^*$ is a normalized $\Delta_m S$ (per residue or per gram) at $T = T_S^*$. The integration constant $\Delta_m S^*$ is a constant with respect to $T$ and, thus, might *a priori* depends on each protein, *i.e.* on each $\Delta C_P$. However, the explicit assumption

[1] It is easy to show that the same results would be obtained by considering first $\Delta_m S$ *vs.* $\Delta C_P$

made in (8, 9) on the universality of the melting of globular proteins implies that the normalized $\Delta_m S^*$ has the same value for all globular proteins. Therefore, within the frame of the latter assumption, equation 2 shows that, indeed, the linearity of $\Delta_m H$ with $\Delta C_P$ implies that of $\Delta_m S$.

**The equality $T_H^* = T_S^*$ is a consequence of thermodynamics and of the assumed universality of protein melting.** Now, the crucial point is that the experimental slope of $\Delta_m S$ *vs.* $\Delta C_P$ must be explained by $Ln\,T/T_S^*$ in equation 2, thus by the value of $T_S^*$. In (8), $T_S^*$ was determined empirically to explain the observed slope $(-0.253 \pm 0.016)$ and it was found that $T_S^* = T_H^*$ was very satisfactory since with $T = 298\,K$ and $T_S^* = T_H^* = (383 \pm 4)K$, the calculated slope is $-0.251 \pm 0.01$. In fact, there is no need of relying on an empirical guess because it may be shown that $T_S^* = T_H^*$ is also imposed by thermodynamic considerations. For that, $\Delta_m G$ is determined by integration of $\partial \Delta_m G/\partial T = -\Delta_m S$ with $\Delta_m S$ from equation 2, which gives :

$$\Delta_m G = \Delta_m G^* + (\Delta C_P - \Delta_m S^*)(T - T_S^*) - \Delta C_P\, T\, Ln \frac{T}{T_S^*} \tag{3}$$

where $\Delta_m G^*$ is the value of the normalized $\Delta_m G$ at $T = T_S^*$. Here also, the argument of universality requires that $\Delta_m G^*$ be the same for all globular proteins. By using $\Delta_m H = \Delta_m G + T\,\Delta_m S$, it is obtained :

$$\Delta_m H = \Delta_m G^* + T_S^*\,\Delta_m S^* + \Delta C_P\,(T - T_S^*) \tag{4}$$

Of course, this must be compatible with equation 1, which implies $\Delta_m H^* = \Delta_m G^* + T_S^*\,\Delta_m S^*$ and $(T - T_H^*) = (T - T_S^*)$. Both consequences require $T_H^* = T_S^*$, which achieves the proof. The controversy on $T_H^* = T_S^*$ (10, 11) is thus resolved.

#### 3 $\Theta_T$ determination and MD considerations

A *Mathematica* notebook illustrating several aspects is available on request.

**The 'simple' and 'generalized' Gamma distributions in use.** In the main text it is explained how Gamma distributions are used to represent the histogram of the variation $\Delta E = E_1 - E_0$ of the potential energy normally obtained after a molecular dynamics (MD) simulation (see section $\Theta_T$ *estimate without continuous X variable*). A 'simple' Gamma distribution $\Gamma_1[\alpha,\beta]$ and a 'generalized' Gamma distribution $\Gamma_2[\alpha,\beta,\gamma,\mu]$ [2] are used according to (12):

$$\Gamma_1[\alpha,\beta](x>0) = \frac{\beta^{-\alpha} x^{\alpha-1} e^{-\frac{x}{\beta}}}{\Gamma(\alpha)}$$
$$\Gamma_2[\alpha,\beta,\gamma,\mu](x>\mu) = \frac{\gamma\, e^{-\left(\frac{x-\mu}{\beta}\right)^{\gamma}} \left(\frac{x-\mu}{\beta}\right)^{\alpha\gamma-1}}{\beta\,\Gamma(\alpha)} \tag{5}$$

[2]The function $\Gamma$ appearing as $\Gamma(\alpha)$ in equations 5 is the standard notation for the factorial function such that $\Gamma(n) = (n-1)!$ for $n \in \mathbb{N}$, whereas $\Gamma_1$ and $\Gamma_2$ are specific notations in this work.

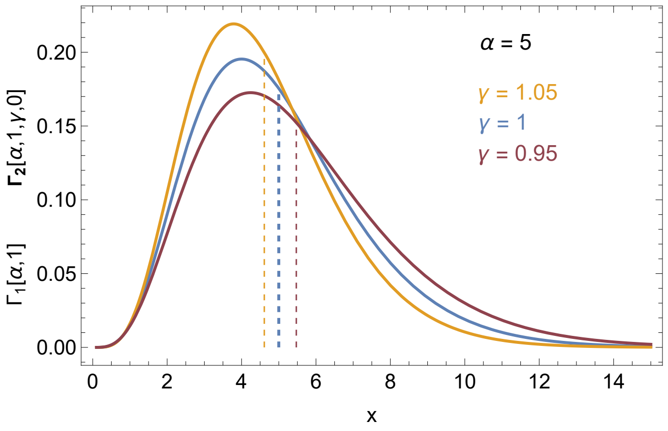


**Fig. S7.** Simple Gamma distribution $\Gamma_1[\alpha,\beta=1](x)$ (blue curve) and generalized Gamma distribution $\Gamma_2[\alpha,\beta=1,\gamma,\mu=0](x)$ for two values of $\gamma \neq 1$. The dashed lines mark the average value $\overline{x}$ of each distribution.

The two distributions are identical for $\gamma = 1$ and $\mu = 0$: $\Gamma_2[\alpha,\beta,\gamma=1,\mu=0] \equiv \Gamma_1[\alpha,\beta]$. They are illustrated in Fig. S7. In this work, $\beta = 1$ was used without loss of generality (see main text). The parameter $\mu$ provides us with the possibility of slightly shifting the whole distribution curve $\Gamma_2$. The simple distribution $\Gamma_1$ allows obtaining an 'exact' evaluation of $\Delta G$ in closed form (equation (17) in the main text), whereas $\Gamma_2$ only allows estimating $\Delta G$ by numerical integration of:

$$\Delta G = -RT\,\ln \int_0^{\infty} \exp\left(-\frac{x}{\overline{x}}\frac{\overline{\Delta E}}{RT}\right) \Gamma_2[\alpha,1,\gamma,\mu](x)\,dx$$
$$\left(\overline{x} = \mu + \frac{\Gamma(\alpha + 1/\gamma)}{\Gamma(\alpha)}\right) \tag{6}$$

where $\overline{x}$ is the average value of a variable $x$ following the distribution $\Gamma_2$. Derivation of $\Delta G$ with respect to $T$ gives:

$$\Delta S = -\frac{\Delta G}{T} + \frac{\overline{\Delta E}}{T} \exp\left(\frac{\Delta G}{RT}\right) \times$$
$$\int_0^{\infty} \frac{x}{\overline{x}} \exp\left(-\frac{x}{\overline{x}}\frac{\overline{\Delta E}}{RT}\right) \Gamma_2[\alpha,1,\gamma,\mu](x)\,dx \tag{7}$$

which can also be obtained by numerical integration. The numerical integrations were performed with the option `NIntegrate` in *Mathematica*. Finally, $\Delta H$ is derived from equations 6 and 7. One can then obtain $\Theta_T = \Delta\Delta H/\Delta\Delta S$ by considering the values of $\Delta H$, $\Delta S$ determined by the parameters $\alpha$, $\gamma$, $\mu$, $\overline{\Delta E}$, and of $\Delta H + \Delta\Delta H$, $\Delta S + \Delta\Delta S$ determined by the close values $\alpha + \delta\alpha$, $\gamma + \delta\gamma$, $\mu + \delta\mu$, $\overline{\Delta E} + \delta\overline{\Delta E}$. It is explained in the following how these values can be obtained.

**Assessing the practical utility of the generalized Gamma distribution.** In order to assess the ability of $\Gamma_2$ to account for real situations one set of parameter values was determined so that the variation of $\Theta_T/T$ with $T$ is as close as possible to that for the binding of hydrophobic ligands to proteins (see main text and Fig. 3). This was achieved by defining a Sum Of Squares (SOS) of residuals between the values of $\Theta_T/T$ obtained from a set of parameters $\alpha$, $\delta\alpha$, $\gamma$, $\delta\gamma$, $\mu$, $\delta\mu$, $\overline{\Delta E}$, $\delta\overline{\Delta E}$ and the values of $\Theta_T/T$ from equation 11 in the main text. Each squared residual was at a particular temperature and ten equally spaced temperatures between 280 and 370 K were considered in the SOS. This determined a function SOS $\left(\alpha, \delta\alpha, \gamma, \delta\gamma, \mu, \delta\mu, \overline{\Delta E}, \delta\overline{\Delta E}\right)$ which was minimized by systematic search and use of the functionality `FindMinimum` in *Mathematica*.
